# Raman spectroscopy reveals phenotype switches in breast cancer metastasis

**DOI:** 10.1101/2021.06.02.446487

**Authors:** Santosh Kumar Paidi, Joel Rodriguez Troncoso, Mason G. Harper, Zhenhui Liu, Khue G. Nguyen, Sruthi Ravindranathan, Jesse D. Ivers, David A. Zaharoff, Narasimhan Rajaram, Ishan Barman

## Abstract

The accurate analytical characterization of metastatic phenotype at primary tumor diagnosis and its evolution with time are critical for controlling metastatic progression of cancer. Here, we report a label-free optical strategy using Raman spectroscopy and machine learning to identify distinct metastatic phenotypes observed in tumors formed by isogenic murine breast cancer cell lines of progressively increasing metastatic propensities. Our Raman spectra-based random forest analysis provided evidence that machine learning models built on spectral data can allow the accurate identification of metastatic phenotype of independent test tumors. By silencing genes critical for metastasis in highly metastatic cell lines, we showed that the random forest classifiers provided predictions consistent with the observed phenotypic switch of the resultant tumors towards lower metastatic potential. Furthermore, the spectral assessment of lipid and collagen content of these tumors was consistent with the observed phenotypic switch. Overall, our findings indicate that Raman spectroscopy may offer a novel strategy to evaluate metastatic risk during primary tumor biopsies in clinical patients.

## Introduction

Metastasis to distant organs is the major cause for breast cancer-related mortality [1]. The ability to assess metastatic propensity during the diagnosis of solid primary tumors is critical to arresting future metastatic growth in at risk patients and to reducing overtreatment of patients not at risk to potentially harmful and costly therapies. However, the accurate determination of metastatic tumor phenotype is particularly challenging due to the involvement of multiple players beyond the cancer cells, such as the components of tumor microenvironment and the composition of secondary sites (pre-metastatic niches) [2–5]. The existing methods provide limited objective insights into the metastatic risk of solid tumors at the primary diagnosis. While emerging research is focused on leveraging the knowledge of mutational landscape of primary tumors as well as biological and physical characterization of circulating tumor cells (CTC) in vitro (e.g., microfluidic systems) and circulating tumor DNA (ctDNA), the clinical adoptions of these methods are either hindered by the requirement of labor-intensive processing or restricted by the knowledge of limited markers of metastatic progression [6–15]. Therefore, there is an urgent need for novel analytical technologies that can provide direct readout of the metastatic potential from the solid tumor samples with minimal perturbation.

Optical spectroscopy and imaging have emerged as attractive platforms to probe biomolecular composition and metabolic status of tumors and their stroma [16–18]. Raman spectroscopy, a non-invasive method based on inelastic scattering of light, is particularly attractive for label-free quantitative tissue analysis due to the adequate penetration depth of near-infrared (NIR) light in tissue and a lack of spectral interference from water content [19–21]. Raman spectroscopy and imaging have been employed previously to study a range of biomedical problems [20–22] including primary tumor composition [23–26], breast cancer microcalcifications [25, 27], formation of pre-metastatic niches [28], response to therapy [29–31], and single-cell phenotyping [32]. However, a systematic study of metastatic phenotypes associated with tumors derived from isogenic cancer cells of progressively increasing metastatic abilities is currently lacking. Additionally, the sensitivity of spectral markers derived from optical spectroscopic measurements to identify tumor phenotypes that result from subtle alterations in expression of genes implicated in metastasis has never been tested.

In this report, we evaluate the ability of Raman spectroscopy to identify distinct phenotypes associated with metastatic propensity by employing an isogenic panel of murine breast cancer cell lines – 4T1, 4T07, 168FARN, 67NR – where each of the cell lines are only capable of accomplishing specific steps in the metastatic process [33] (**Fig. 1**). We use multivariate curve resolution-alternating least squares (MCR-ALS) decomposition of spectra and supervised classification analysis using random forests to reveal putative molecular markers of progression and predict the metastatic phenotype in tumors. We also used gene deletion/knockdown variants of the 4T1 cell line, which has the highest metastatic potential, to determine if targeting metastasis-promoting genes (TWIST1, FOXC2, CXCR3) would cause tumors grown from these cell lines to be classified differently. Prior research has shown that silencing of TWIST1, FOXC2, and CXCR3 genes individually in the aggressive 4T1 cell line results in significant loss of metastatic abilities [34–36]. We show that tumors grown from 4T1 cells with these genes silenced are classified as having lower metastatic potential. We also demonstrate the ability of Raman spectroscopy to delineate differences between the phenotype switches achieved via different biological pathways mediated by these gene silencing experiments.

**Figure 1.**
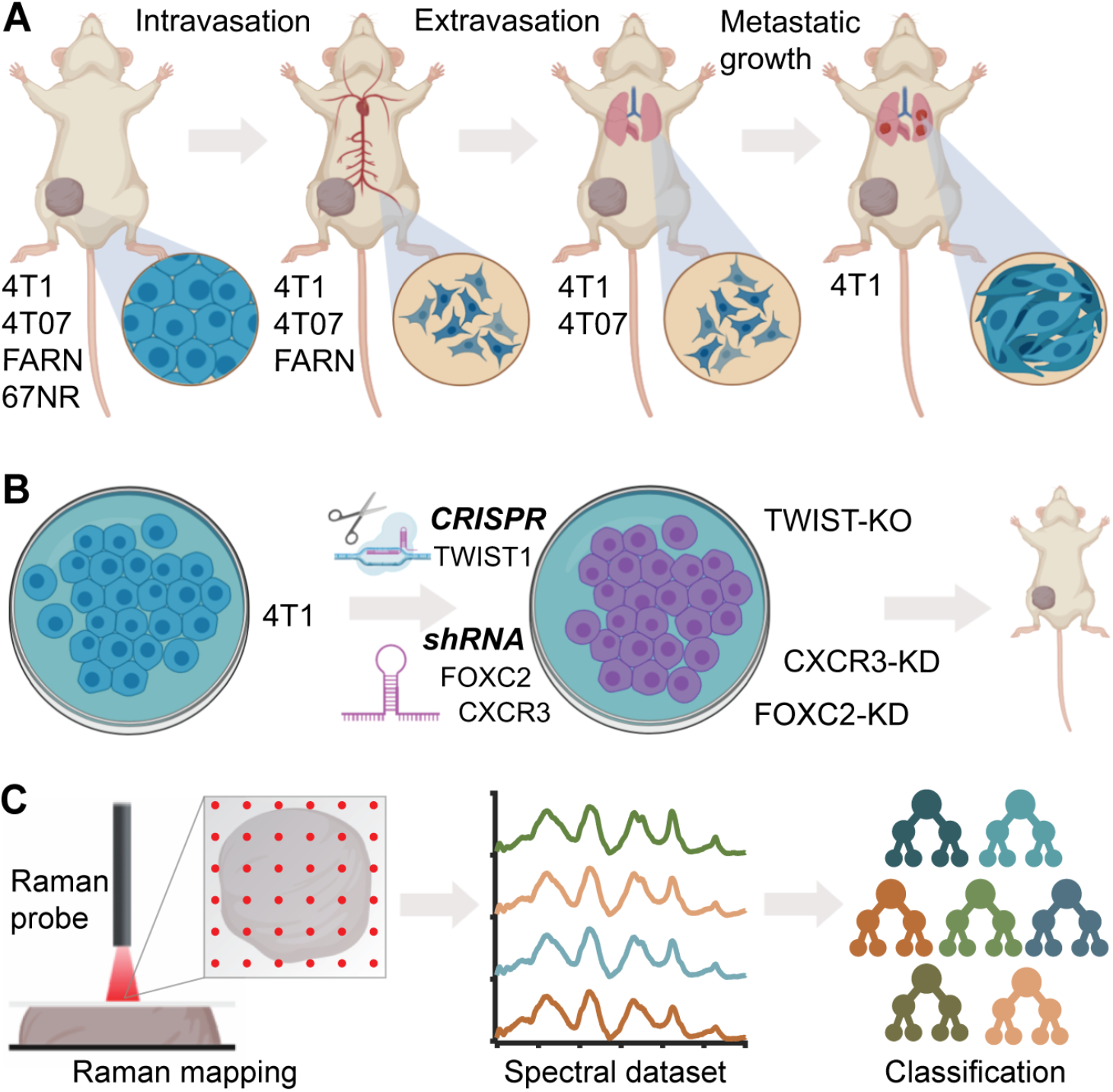
Label-free Raman spectroscopy for identifying metastatic phenotypes. (**A**) The different steps of metastatic cascade accomplished by the tumors formed by each of the cell lines of the 4T1 tumor model employed in this study are shown. (**B**) The tumors formed by the 4T1 cells silenced for the expression of metastasis-promoting genes are employed for spectroscopic measurements. (**C**) The overview of Raman mapping of the tumors and spectral analysis is presented.

## Results

### Raman spectroscopy reveals differences between isogenic breast tumors of varying metastatic potential

We used the mouse mammary tumor model comprised of four isogenic cell lines of varying metastatic potential – 67NR, 168FARN (FARN), 4T07, and 4T1 – that were originally derived from a single mammary tumor in a wild-type BALB/c mouse [33]. Each of these cell lines form robust primary tumors in BALB/c mice but exhibit progressive differences in their metastatic abilities (**Fig. 1A**) [34]. The non-metastatic 67NR cells fail to intravasate into the circulation. Of the remaining three cell lines, FARN cells fail to extravasate into the lungs. While both 4T07 and 4T1 cells extravasate into the lungs, only 4T1 cells manage to establish visible metastatic nodules and 4T07 cells have been shown to disappear from lungs following removal of the primary tumor. Therefore, these four cell lines form an excellent model to study distinct steps of cancer metastasis. In the current study, we employed tumors formed by 67NR (n=8), FARN (n=6), 4T07 (n=5), and 4T1 (n=8) cells in BALB/c mice. Our spectral dataset comprised of 6362 spectra (average *ca*. 235 spectra per tumor) collected from these 27 tumors (**Fig. 2A**). In addition, the spectral dataset also included 8589 spectra from 34 tumors derived by using the variants of 4T1 cells with specific genetic modifications (described later).

**Figure 2.**
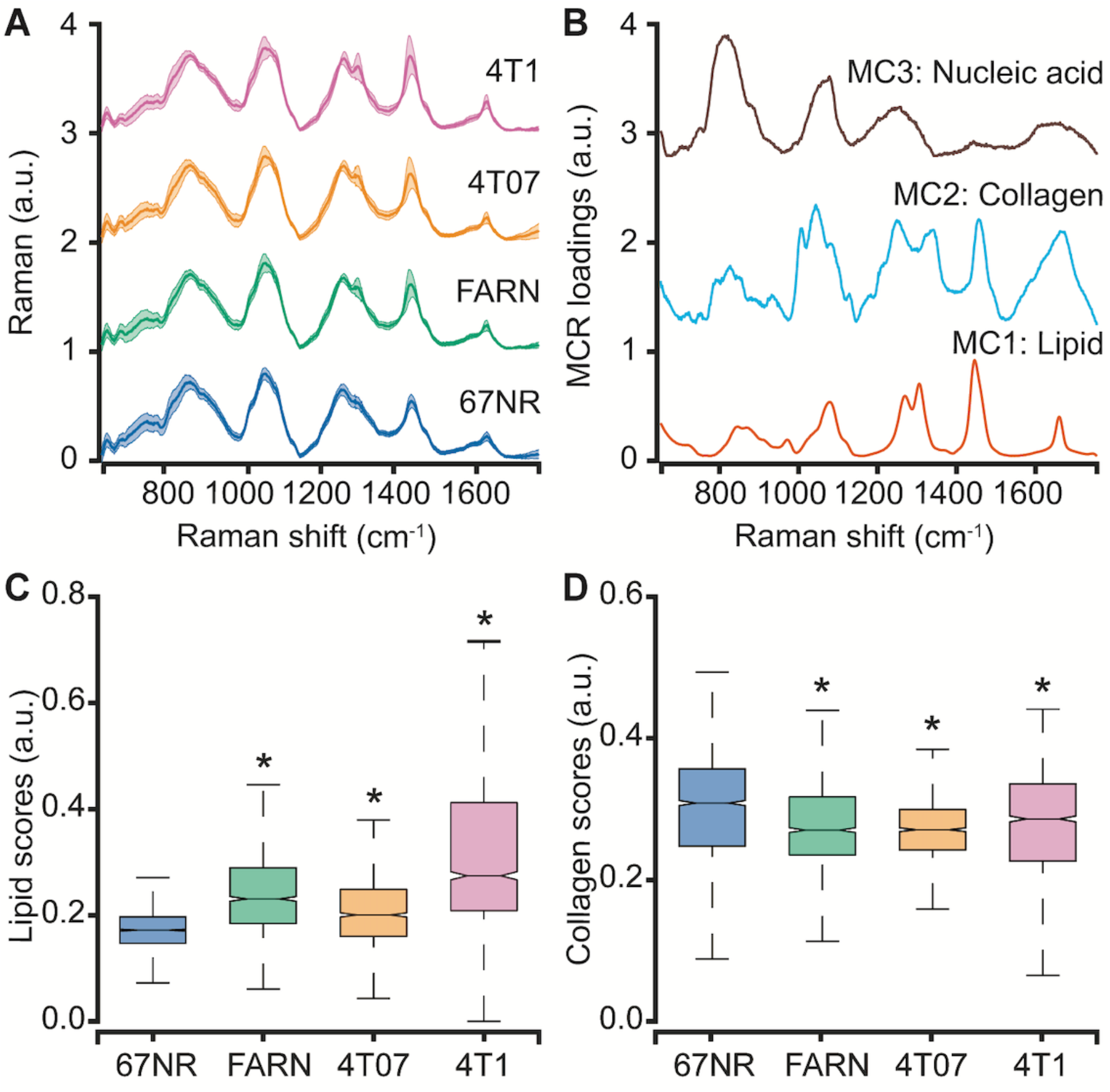
Spectral differences between tumors of varying metastatic potential. (**A**) The mean (dark line) and 1 standard deviation (shaded region) of the Raman spectra collected from isogenic tumors of varying metastatic potential are shown. (**B**) A subset of constituent spectra derived using MCR-ALS decomposition of Raman spectral dataset that harbor features of lipids, collagen and nucleic acids are plotted. The box and whisker plots show the variation of scores of (**C**) lipid-like and (**D**) collagen-like MCR-ALS components with metastatic potential of the tumors. Statistical significance as assessed by Wilcoxon rank-sum test p-value < 0.05 for each metastatic tumor group in comparison with non-metastatic 67NR group are denoted using asterisks.

We subjected the entire spectral dataset to multivariate curve resolution – alternating least squares decomposition (MCR-ALS) for dimensionality reduction and identification of spectral constituents [37]. Due to the positivity constraints imposed on the derived component spectra and their scores, MCR-ALS decomposition has been shown to provide components that resemble pure constituents of the specimen without prior knowledge of its composition. For the entire spectral dataset in the present study, a five-component MCR-ALS decomposition (**Fig. 2B**) provided components that resemble lipid (MC1), collagen (MC2), and nucleic acid (MC3) based on their characteristic spectral features detailed in **Table ST1** (**Supporting Information**). The remaining two components, respectively, captured contributions due to formalin contamination and mixed spectral features (**Fig. S1, Supporting Information**). We compared the distribution of scores of each MCR-ALS component across the four tumor groups of varying metastatic potential. We observed that the median scores of the lipid-like component (MC1) increased significantly for the metastatic tumors formed by FARN, 4T07, and 4T1 compared with the non-metastatic 67NR tumors and that the increase was highest for the 4T1 tumors (**Fig. 2C**). Similarly, we found the median MCR-ALS scores for the collagen-like components decreased for the tumors in FARN, 4T07, and 4T1 groups in comparison with 67NR tumors (**Fig. 2D**). While these component scores show significant differences between the non-metastatic 67NR tumors and the remaining tumor groups that show progressively higher metastatic abilities, they do not vary uniformly across the classes. Therefore, the univariate analysis based on the component scores alone is not sufficient to predict the metastatic potential of the tumors.

### Supervised classification using random forests predicts stage-specific metastatic phenotypes

We used supervised classification based on random forests with a leave-one-mouse-out approach to train classifier models on the spectral dataset and predict the metastatic phenotype of the tumors. First, we trained and tested multi-class random forest classifiers on spectra from all the four tumor groups (**Fig. 3A**). The leave-one-mouse-out strategy involves training the classifier on spectral dataset by excluding all the spectra from one mouse in each iteration and subjecting them as an independent test dataset to the developed model. By eliminating the representation of spectra from test mice in the training dataset, this approach prevents overfitting of the classifier and provides robust models that incorporate biological variability across the tumors obtained from different mice. The predicted class label is determined for each test mouse based on the majority voting of the spectral predictions. We observed that the leave-one-mouse-out approach provided accurate predictions for most mice in the non-metastatic 67NR (7/8) and highly metastatic 4T1 tumors (8/8). The tumors in the intermediate groups FARN (4/6) and 4T07 (4/6) were either accurately classified or misclassified into their neighboring groups in terms of metastatic potential. Together, these observations show that random forest classifiers are not only able to accurately distinguish between the non-metastatic and highly metastatic phenotypes, but also satisfactorily estimate the intermediate phenotypes characterized by relatively subtle differences in the spectrum of metastatic progression. The classification accuracy, particularly of the intermediate groups (FARN and 4T07), should be further boosted with the inclusion of more mice in the dataset.

**Figure 3.**
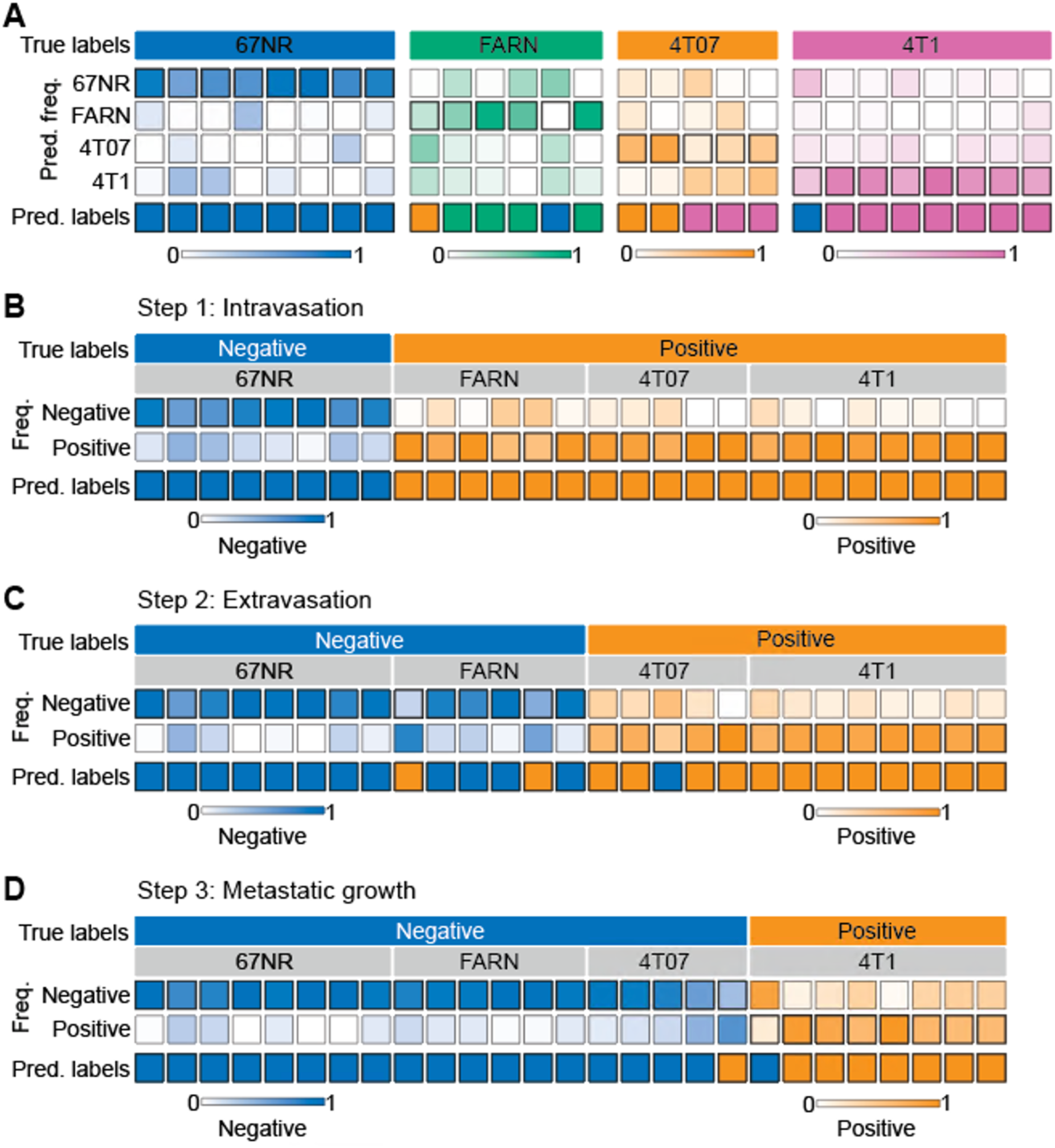
Supervised classification of metastatic potential and stage-specific phenotypes. The results for leave-one-mouse-out random forest classification are shown as heatmaps for the prediction of – metastatic phenotype of the tumors formed by the four cell lines with differential metastatic potential (**A**) and stage specific metastasis abilities to accomplish intravasation (**B**), extravasation (**C**), and metastatic growth (**D**). The true labels for the analysis (negative or positive for each step) in panels **C-D** are assigned based on known behaviors of these tumors in vivo. The top and bottom rows in each heatmap, respectively, show the true labels and predicted labels of the individual mice (columns) in each group. The labeled central rows in the heatmaps show the distribution of the predicted labels for spectra from each mouse into the classes in the training dataset. The overall class prediction for each mouse is obtained by thresholding on the prediction frequencies.

Next, we sought to determine if the random forest classifiers derived from the Raman spectra can specifically distinguish tumors that complete specific steps of the metastatic cascade from those that do not. We trained and tested binary random forest classifiers with the leave-one-mouse-out analysis for intravasation, extravasation, and metastatic growth. For intravasation, we labeled spectra from the 67NR tumors as intravasation-negative and the spectra from the remaining tumors as intravasation-positive and subjected them to binary leave-one-mouse-out random forest classification (**Fig. 3B**). We observed that all tumors were classified accurately as belonging to either negative or positive classes. Using a similar approach for extravasation by labeling 67NR and FARN tumors negative and 4T07 and 4T1 tumors positive, we noted that only two FARN tumors and one 4T07 tumor were misclassified as positive and negative respectively for extravasation (**Fig. 3C**). All remaining tumors were classified accurately according to their extravasation capacity. Finally, we tested the random forest classifiers for overt metastatic growth. We labeled the spectra from the 67NR, FARN and 4T07 tumors as negative and spectra from 4T1 tumors as positive for metastatic growth (**Fig. 3D**). We found that only one tumor each of the 4T07 and 4T1 groups were respectively misclassified as positive and negative for metastatic growth. The accurate prediction of metastatic propensity and identification of specific steps in the metastatic cascade show that Raman spectra can capture subtle biomolecular features of the tumor and its microenvironment that define its metastatic outcome.

### Random forest analysis reveals stage-specific Raman spectral markers of metastatic progression

Random forests provide intrinsic capacity to rank spectral features based on their contribution to the prediction accuracy. We performed random forest classification on the entire spectral dataset for each step of metastasis and determined the top five spectral features that contribute the most to the classification accuracy. For intravasation, we found that the classification is governed by spectral features at 1042 cm^−1^, 1213 cm^−1^, 1330 cm^−1^, 1432 cm^−1^, and 1566 cm^−1^ (**Fig. 4A**). We observed that the classification for extravasation is governed by a distinct set of spectral makers at 670 cm^−1^, 962 cm^−1^, 1372 cm^−1^, and 1664 cm^−1^ in addition to the overlap with the intravasation markers at 1213 cm^−1^ (**Fig. 4B**). Similarly, the classification of the spectra based on the metastatic growth step were governed by features at 862 cm^−1^, 1098 cm^−1^, 1254 cm^−1^, 1462 cm^−1^, and an overlapping feature with extravasation set at 1664 cm^−1^ (**Fig. 4C**). We plotted these top five discriminating wavenumbers for each step as a Venn diagram (**Fig. 4D**) and observed overlap in the sets of spectral markers only between the neighboring steps of the metastatic cascade (between intravasation and extravasation sets as well as extravasation and metastatic growth sets). The band assignments for the identified predictors for all the steps are tabulated in **Table ST2** (**Supporting Information**). Together with the lack of overlap between intravasation and metastatic growth sets, the random forest derived spectral markers provide evidence that distinct metastatic abilities are governed by different biomolecular constituents of the tumors and that Raman spectroscopy can delineate the differences and similarities in their spectral signatures.

**Figure 4.**
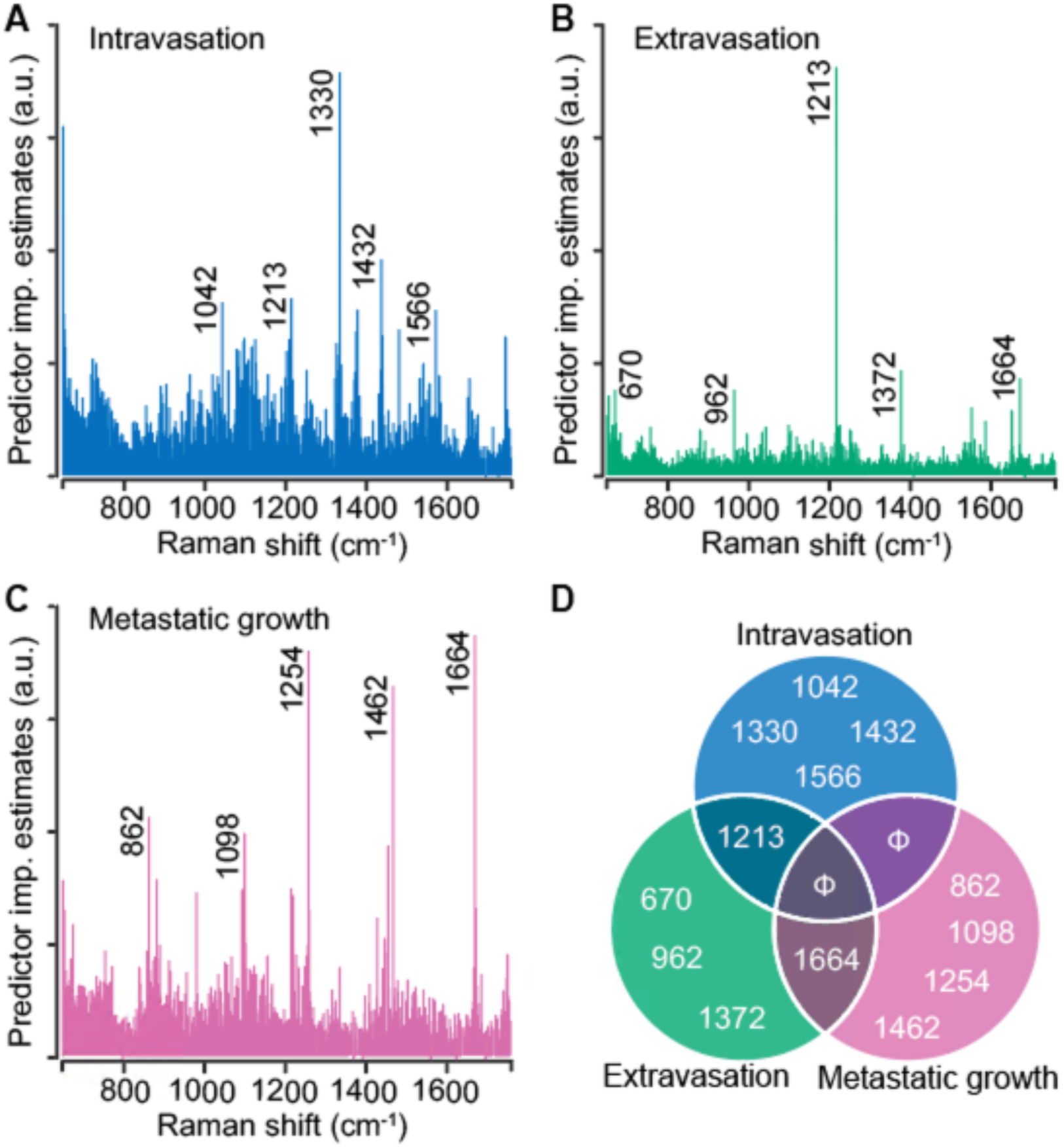
Stage-specific spectral markers of metastatic progression. The predictor importance estimates derived from random forest classification of Raman spectra based on their stage-specific metastatic abilities are shown for intravasation (**A**), extravasation (**B**), and metastatic growth (**C**). The five most important (non-neighboring) Raman features (cm^−1^) are highlighted on these plots and presented as a Venn diagram (**D**) to visualize the overlap between different steps.

### Raman spectroscopy allows identification of metastatic phenotype of tumors formed by silencing genes that drive metastatic progression in 4T1 cells

We used 4T1 cells after CRISPR/Cas9 knockout of TWIST1 gene (TWIST-KO, n=8) and shRNA knockdown of FOXC2 and CXCR3 genes (FOXC2-KD, n=7 and CXCR3-KD, n=6) to grow tumors in BALB/c mice and determine the accuracy of Raman spectroscopy in identifying the metastatic potential of the resultant tumors. We also used the vector control cells for the shRNA knockdowns to grow respective control tumors (FOXC2-VC, n=7 and CXCR3-VC, n=6). Prior studies have shown that silencing of TWIST1, FOXC2, and CXCR3 genes in 4T1 cells resulted in a reduction of the metastatic potential of the resultant tumors in mice and a lower lung metastatic burden [34–36]. To test if the Raman spectra of the tumors derived from the silenced cells capture these phenotypic changes, we subjected the spectra acquired from these tumors as test datasets to the random forest classifier models trained on the spectra from the four original tumors classes – 67NR, FARN, 4T07, and 4T1 (**Fig. 5A**). The per-mouse predictions for the TWIST-KO tumors revealed that a majority (6/8) of them were classified as non-metastatic 67NR tumors and neither were classified as tumors derived from 4T1 cells, from which the silenced cells were derived. Next, we inspected the per-mouse predictions for FOXC2-KD tumors and compared them with their vector control FOXC2-VC tumors. We observed that most of the FOXC2-KD tumors were classified as the lower metastatic FARN (3/7) or 4T07 (2/7) tumors than the parental 4T1 tumors (2/7). In the vector control FOXC2-VC tumors, however, we found that most tumors were classified as 4T1 (4/7) and 4T07 (2/7) tumors. Similarly, we noted that the CXCR3-KD tumors were classified largely as FARN (2/6) and 4T07 (3/6) tumors whereas the vector control CXCR3-VC tumors were predominantly classified as the parental 4T1 (4/6) tumors. Our results, here, provide the first evidence that optical spectroscopic signatures can potentially capture functional differences in the tumor states that arise from alterations in gene expression at cellular level.

**Figure 5.**
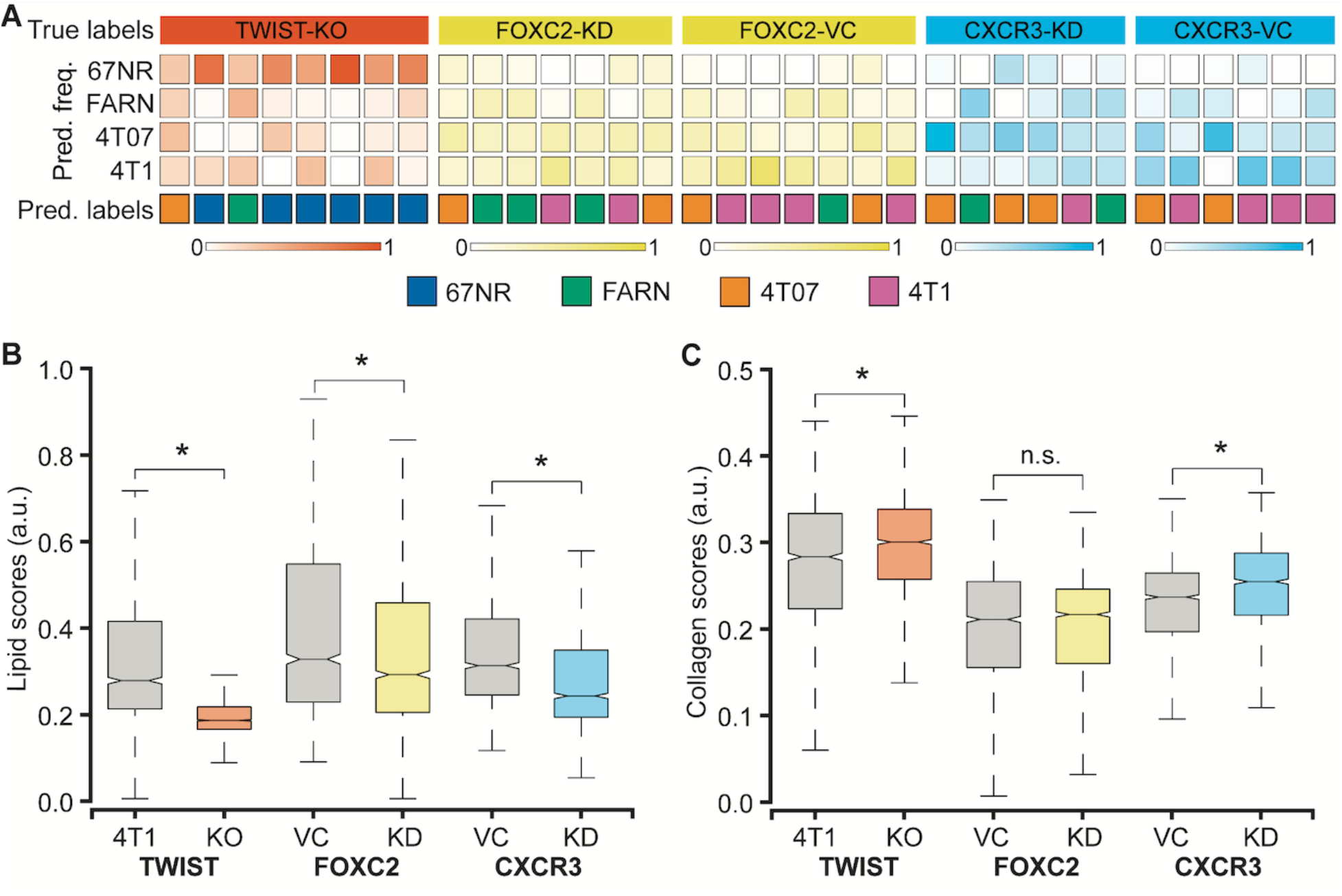
Raman spectroscopic identification of metastatic phenotypes due to subtle alterations in gene expression. (**A**) Heatmap representation of random forest classification of tumors from the three 4T1-variant cell lines (knockdown and control). The classifier model was trained on data from the original tumor panel - 67NR, 168FARN, 4T07, and 4T1. The overall class prediction in the bottom row for each mouse is obtained by thresholding on the prediction frequencies of the four classes in the labeled intermediate rows. A comparison of the MCR-ALS scores between the knockdown and control tumors for the components resembling lipids (**B**) and collagen (**C**) is shown using box and whisker plots. Statistical significance as assessed by Wilcoxon rank-sum test p-value < 0.05 for each genetically altered tumor group in comparison with their respective control groups is denoted using asterisks.

To further verify that the metastatic phenotype identification of the tumors is driven by the changes in the tumor microenvironment that result from alterations in gene expression, we compared the MCR-ALS scores of the Raman spectra obtained from the TWIST-KO, FOXC2-KD, and CXCR3-KD tumors with their respective controls. We found a significant reduction in the scores of the lipid-like component (MC1) in the TWIST-KO tumors in compared to their control 4T1 tumors (**Fig. 5B**). Similar decreases in the scores of the lipid-like component were observed for the FOXC2-KD and CXCR3-KD tumors when compared to their respective vector controls FOXC2-VC and CXCR3-VC. For the collagen-like component (MC2), we found that the scores of the TWIST-KO and CXCR3-KD tumors increased significantly when compared to their respective controls (**Fig. 5C**). The decrease and increase, respectively, in scores of the lipid-like and collagen-like components due to the knockdown of genes essential for metastasis in the highly metastatic 4T1 cells are consistent with the observed differences between non-metastatic 67NR and variably metastatic FARN, 4T07, and 4T1 tumors in **Figure 2**. Together with the random forest classification results, the consistent changes in the MCR-ALS scores of the pure component-like constituents improve our confidence in the ability of Raman spectroscopy to identify biomolecular differences in the tumors that result from a phenotypic switch due to alterations in gene expression.

### Raman spectroscopy can distinguish between similar metastatic phenotype that result from distinct alterations in gene expression

Prior microarray analysis of the tumors formed by 67NR, FARN, 4T07, and 4T1 cell lines has identified key genes are important drivers of each step of the metastatic cascade [34]. Weinberg and coworkers have performed the same stage-specific comparisons of the microarray data obtained from the four tumors and identified unique sets of genes that are most significantly upregulated or downregulated in groups that show differential intravasation, extravasation and metastatic growth [34]. Their gene expression analysis showed that TWIST1 and CXCR3 genes are overexpressed in tumors formed by cell lines capable of intravasation - FARN, 4T07, and 4T1 – compared with the non-metastatic 67NR cells. Similar analysis showed that FOXC2 is overexpressed in the highly metastatic 4T1 tumors compared with the 67NR, FARN, and 4T07 tumors. In addition to these genes, the microarray analysis revealed several other genes that are overexpressed in tumors with distinct metastatic capabilities. Based on these results, we selected 24 genes overexpressed in metastatic tumors to build a protein-protein interaction network and explore their roles in different biological processes responsible for cancer metastasis using the STRING-11 (http://string-db.org) analysis software. The network in **Figure 6A** shows the protein-protein interactions between the overexpressed genes and a subset of biological processes that are governed by them. We found evidence of co-expression of TWIST1 and FOXC2 as well as evidence of experimentally determined relationship between TWIST1 and CXCR3 via MMP9. We also found that the three genes contributed to overlapping, yet different, biological processes involved in cancer metastasis.

**Figure 6.**
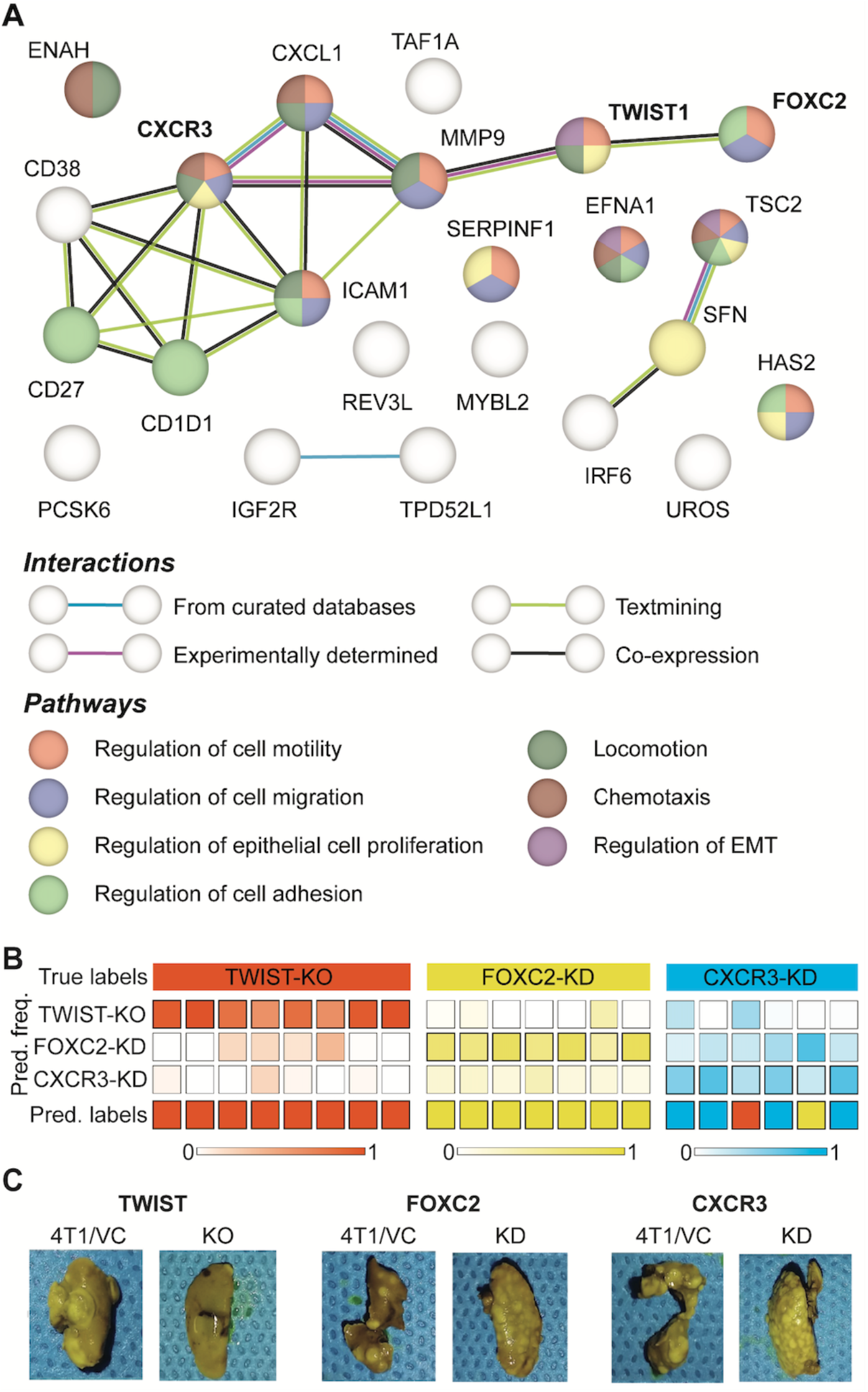
Differences between metastatic phenotype switches due to distinct alterations in gene expression. (**A**) Protein-protein interaction network of genes identified as overexpressed in the tumors accomplishing intravasation, extravasation, metastatic growth is shown. The network nodes are colored by their pathway membership and the interactions between nodes are colored by type, as listed in the respective legends. (**B**) The results of leave-one-mouse-out random forest classification of the spectra from tumors obtained by silencing TWIST1, FOXC2, and CXCR3 expression in 4T1 cells are shown. The overall class prediction in the bottom row for each mouse is obtained by thresholding on the prediction frequencies of the three classes in the labeled intermediate rows. (**C**) Representative photographs of metastatic lungs of mice harboring tumors obtained by silencing TWIST1, FOXC2, and CXCR3 expression in 4T1 cells are shown along with their corresponding controls.

To test if the differential regulation of breast cancer metastasis due to TWIST1, FOXC2, and CXCR3 genes results in spectroscopically distinct phenotypes, we performed a three-class random forest classification using the leave-one-mouse-out routine with spectral data from just the three altered tumor groups (**Fig. 6B**). While all the TWIST-KO and FOXC2-KD tumors were classified accurately, two CXCR3-KD tumors were misclassified as TWIST-KO and FOXC2-KD tumors, respectively. The overall high accuracy of the three-class classifier shows that Raman spectroscopy has the potential to distinguish between similar phenotypic states achieved via different biological pathways. These differences were further corroborated by the assessment of macrometastatic nodules (**Fig. 6C**) in lungs of the mice bearing tumors grown from the cell lines silenced for the expression of TWIST1, FOXC2, and CXCR3, that showed distinct differences in the metastatic burden. The significantly larger reduction in the metastatic burden observed in TWIST-KO group compared to the FOXC2-KD and CXCR3-KD groups and the relative misclassification of the CXCR3-KD spectra could also be partially attributed to the relatively more robust silencing of TWIST1 due to CRISPR/Cas9 knockout at the genome level in comparison of shRNA knockdown of FOXC2 and CXCR3 at the mRNA level.

## Discussion

Clinical intervention of cancer metastasis and choice of treatment are dependent on our ability to recognize the metastatic phenotype of primary tumors at initial diagnosis. While prior optical spectroscopic studies of benign and malignant tumors have attempted to assess metastatic potential in spectral terms, the differences in genetic backgrounds of the patient-derived samples limit our ability to delineate the spectral features associated with metastatic phenotypic differences from inter-patient heterogeneity [25, 38]. In the current study, we employed a panel of isogenic breast cancer cells of known but progressively increasing metastatic propensities to grow tumors in mice for molecular phenotyping using Raman spectroscopy [33]. The 4T1 mammary tumor model was employed in the current study due to its ability to spontaneously metastasize to lungs and recapitulate the key steps of metastatic progression in human patients [34]. The MCR-decomposition of spectral dataset from 67NR, FARN, 4T07, and 4T1 tumors revealed significant differences in lipid and collagen content of the non-metastatic 67NR tumors compared to the tumors in the remaining groups (**Fig. 2**). The observed higher lipid content (**Fig. 2C**) in metastatic tumors is in agreement with emerging evidence of an important role of lipid metabolism in cancer metastasis [39, 40]. A recent study by Pascual *et al*. showed that metastasis-initiating cells rely on dietary lipids and express high levels of fatty acid receptor CD36 and genes necessary for lipid metabolism [41]. Similarly, the lower collagen content observed in the metastatic tumors (**Fig. 2D**) could potentially be attributed to the extracellular matrix remodeling due to collagen degradation by stromal cell-derived matrix metalloproteinases [42]. More detailed investigations are necessary to understand the roles of lipid and collagen content in determining the metastatic phenotype of breast tumors due to the multiple roles of these molecules at each step in primary tumor growth and metastatic progression. The lipid and collagen scores, however, did not show monotonic trends with the increasing metastatic potential of the tumor groups. The deviation from monotonic increase for the scores of lipid-like MCR component (**Fig. 2C**) resulted from a jump observed for the FARN tumors. Similar deviation from the monotonic decrease of the collagen-like component scores (**Fig. 2D**) has been observed due to an anomalous increase for 4T1 tumors at the end of the spectrum. These deviations, however, do not undermine the significance of the MCR-ALS derived putative markers of metastatic progression.

To utilize the latent information from the entire Raman spectra, we employed random forest classification analyses using a leave-one-mouse-out approach, which permits the treatment of each tumor as an independent test sample against the classifiers trained on the spectral data from the remaining tumors. The near-perfect classification of the tumors in the 4T1 and 67NR groups at both the ends of metastatic propensity (**Fig. 3A**), and the classification of all the misclassified tumors in the intermediate metastatic groups – 4T07 and FARN – exclusively into their neighboring classes demonstrate the power of Raman spectroscopy and random forest classifiers in identifying subtle differences in metastatic phenotypes. It is also important to verify if the classifiers can identify differences in tumors derived from cell lines lacking gene expression associated with critical steps of the metastatic process. The high prediction accuracy obtained for the leave-one-out random forest classifiers trained on the outcomes for intravasation, extravasation, and metastatic growth (**Fig. 3B-D**) demonstrated the feasibility of using Raman spectroscopy for accurate localization of the primary tumors in the metastatic cascade. Furthermore, the intrinsic feature importance ranking provided by random forests showed that the stage-specific identification is driven by distinct sets of spectral markers with minimal overlap instead of a universal set of markers (**Fig. 4**). It is interesting to note that the spectral marker overlap can only be seen between adjacent steps in the metastatic cascade. This observation is consistent with the complexity of metastatic process, which is characterized by various stagespecific molecular biological processes.

The 4T1 family of isogenic mammary tumors has been used by several researchers to identify important genes that drive breast cancer metastasis such as TWIST1, FOXC2, and CXCR3 [34–36]. Inhibition of these genes, via siRNA and shRNA, in 4T1 cells resulted in substantial reduction of metastatic nodules in the lungs. Therefore, these observations provided a rationale for our investigation into the spectral characteristics of tumors formed by 4T1 cells after silencing the expression of TWIST1, FOXC2, and CXCR3 genes. The classification of the majority of tumors silenced for critical genes responsible for metastasis into the lower metastatic groups and their respective controls into the high metastatic 4T1 group (**Fig. 5A**) shows that Raman spectroscopic measurements are capable of capturing the phenotype switches associated with subtle alterations in gene expression. Furthermore, the statistically significant changes in the MCR-ALS scores of lipid-like and collagen-like components (**Fig. 5B-C**) in the direction consistent with their phenotype switch validates the existence of a direct relationship between the spectral markers obtained by MCR-ALS analysis and the target tumor phenotype.

While the silencing of TWIST1, FOXC2, and CXCR3 genes is known to reduce metastatic burden, their contribution to metastasis is driven by different biological pathways. For example, TWIST1 and FOXC2 contribute to metastasis by regulating epithelial to mesenchymal transition (EMT) [35, 43]. However, while TWIST1 regulates EMT by suppressing the expression of epithelial marker E-cadherin, FOXC2 does not alter E-cadherin levels and is responsible for the induction of mesenchymal phenotype [43, 44]. CXCR3, on the other hand, regulates metastasis through impairment of host anti-tumor immunity due to suppression of IFN-γ production and T cell expansion [36]. The differential metastatic burden observed in the lungs of mice bearing TWIST-KO, FOXC2-KD, and CXCR3-KD tumors compared to their controls are in agreement with these prior observations. Furthermore, the accuracy of three-class random forest classifiers trained on the spectra obtained from TWIST-KO, FOXC2-KD, and CXCR3-KD in our current study (**Fig. 6**) hints at the potential of Raman spectroscopy to capture the differences between similar metastatic phenotypes that result from loss of metastatic abilities via distinct biological pathways.

## Conclusion

In summary, our results show that Raman spectroscopy and machine learning can provide a potent combination for identification of differences in metastatic phenotypes that result from subtle alterations in gene expression. The rich molecular information captured by the spectra can be leveraged to determine important markers of metastatic progression. Our strategy coupled the uniform genetic background provided by the isogenic mouse model of breast cancer metastatic progression with gene silencing for known mediators of metastasis in this model. This allowed us to attribute the observed differences in the spectral patterns to distinct steps in the metastatic cascade. We envision that this approach will be adopted to study other disease systems where phenotypic differences are guided by subtle alterations in gene expression. Future studies will leverage the molecular specificity of Raman spectroscopy for monitoring the evolution of metastatic risk in response to primary tumor therapy by inspecting pre-treatment and post-treatment biopsies. In addition, we plan to build on the current results to develop new label-free analytical solutions for identifying the best gene targets to assist the development of novel cancer gene therapeutics.

## Material and Methods

### Cell culture and tumor xenografts

The cell lines, 67NR, 168FARN, 4T07, and 4T1 were originally derived from a spontaneous breast tumor growing in a Balb/c mouse and were kindly provided by Dr. Fred Miller (Karmanos Cancer Institute, Detroit, MI) [33]. The TWIST-KO cells were generated by deleting the TWIST1 gene in 4T1 cells using CRISPR/Cas9 system. The 20-base pair sgRNA targeting the TWIST gene (5’-TTGCTCAGGCTGTCGTCGGC-3’) was identified using the sgRNA guide tool by Zhang laboratory at MIT (Cambridge, MA). The sgRNA was cloned into pCasGuide-EF1a-GFP plasmids by OriGene (Rockville, MD), which were expanded in E.Coli bacteria and isolated using the QIAGEN Plasmid Maxi Kit. The 4T1 cells were seeded in a 6-well plate at a density of one million cells per well and 10 μg of plasmids in lipofectamine 3000 were added for transfection and verified using a Nikon TiE fluorescence microscopy after 24 to 48 hours. The transfected 4T1 cells suspended in phosphate buffered saline (PBS) were filtered through a 50 μm filter into a FACS tube for sorting based on GFP expression through BD Biosciences (San Jose, CA) Aria III FACS system. The 5% of transfected cells with the highest GFP expression were selected and incubated for 7 to 14 days. From the 13 clones obtained from the cell colonies, the clone expressing least TWIST1 gene were selected. The previously characterized FOXC2-KD shRNA knockdown clones and their vector controls FOXC2-VC clones derived from 4T1 cells were obtained from Dr. Sendurai A. Mani (MD Anderson Cancer Center, Houston, TX) [35, 44]. The previously characterized CXCR3-KD shRNA knockdown clones and their vector controls CXCR3-VC clones derived from 4T1 cells were obtained from Dr. Li Yang (National Cancer Institute, Bethesda, MD) [36]. The cells were cultured in Dulbecco’s Modified Eagle’s Medium (DMEM) with the addition of 10% (v/v) fetal bovine serum (FBS), 2 mM L-glutamine, 1% (v/v) nonessential amino acids, and 1% (v/v) penicillin-streptomycin and maintained in a humidified incubator at 5% CO_2_ and 37°C. For each cell line, about 150,000 to 250,000 cells (4 million cells for the 168FARN cells) suspended in 100 μl of saline were injected into the flanks of Balb/c mice to grow xenografts. Tumors were excised when they reached an average volume of 200 mm^3^ or if they started to show signs of necrosis. The snap-frozen lungs for metastasis assessment were thawed by placing them in 20-30ml of PBS at 4°C for 15 minutes at room temperature. To remove the OCT surrounding the lung tissue, the submerged samples were occasionally agitated at 10 rpm. Subsequently, each lung was placed in a centrifuge tube in 5mL of Bouin’s Solution at room temperature and incubated for 3 days. All experiments were approved by the Institutional Animal Care and Use Committee at the University of Arkansas (IACUC protocol 18062).

### Raman spectroscopy

The excised tumors were snap-frozen in liquid nitrogen and stored at −80°C. The frozen tumors were thawed in PBS and fixed in 10% neutral buffered formalin prior to the Raman measurements. The formalin fixed tumors were washed in PBS and sandwiched between a quartz coverslip and an aluminum plate for Raman mapping measurements. A previously described fiber probe-based portable clinical Raman spectroscopy system was used to perform Raman mapping in the current study [45]. Briefly, the system is comprised of an 830 nm diode laser (Process Instruments, maximum power: 500 mW) for excitation, a spectrograph (Holospec f/1.8i, Kaiser Optical Systems), and a thermoelectrically cooled CCD camera (PIXIS 400BR, Princeton Instruments). The laser power of ca. 20 mW was delivered to the tissue via a fiber-optic probe mounted on a motorized 2-D translational stage (T-LS13M, Zaber Technologies Inc., travel range: 13 mm) to acquire spectra from distinct points on the flattened tumors approximately 1 mm apart. Each spectrum from spatially distinct points was acquired for 5 seconds (10 accumulations of 0.5 seconds each to prevent CCD saturation).

### Data analysis

All Raman spectroscopic data analyses were performed in MATLAB (Mathworks, Natick, MA) environment. The wavenumber axis for the Raman spectra was calibrated using the known features of acetaminophen spectra. The spectra in the 600-1800 cm^−1^ fingerprint region were used for analysis. The spectra were subjected to background removal by iterative fitting and subtraction of a fifth order polynomial and median filtering. Finally, the spectra were vector normalized to ensure that the Euclidean norm of each spectrum is unity.

Multivariate curve resolution-alternating least squares (MCR-ALS) analysis was used to identify pure component-like constituent spectra by reducing the dimensionality of the dataset [37]. MCR-ALS decomposition produces the component loading spectra and their scores for each spectrum in the dataset under positivity constraints on both without necessitating prior knowledge of mixture composition. The MCR-ALS derived scores, which provide surrogates for concentrations of the pure-components, were plotted using box and whisker plots to understand the evolution of each component concentration with the metastatic potential. The few outliers outside the whiskers were not plotted for improved visualization. The statistical significance of the differences in the medians of the MCR-ALS scores across various tumor groups were determined using a two-sided Wilcoxon rank-sum test. The differences in medians were considered significant at a p < 0.05 level.

We employed random forests for supervised classification in this study. The TreeBagger class in MATLAB was implemented with 100 trees to invoke Breiman’s original algorithm [46]. We used a leave-one-mouse-out training strategy to prevent representation of test mice in the training dataset. The random forests classifiers are trained on the entire spectral dataset by excluding all the spectra from one mouse at a time and the obtained models are used to test the spectra of the left-out mouse. The majority predicted label among all the spectra determines the final predicted label for the test mouse. For each left out mouse, 100 iterations of training were performed by random equalization of the study classes in the training set to ensure equal representation. By leveraging the intrinsic capacity of random forests to rank features in the order of their contribution to the prediction accuracy, the predictor importance estimates were derived by training and testing random forest model on the entire spectral dataset using the OOBPredictorImportance argument in the TreeBagger class.

### Protein-protein interaction network

The prominent genes previously identified to be overexpressed in the tumors derived from 4T1 family of cell lines that accomplished each step of metastasis were used to visualize proteinprotein interactions using the STRING-11 (http://string-db.org) analysis software with a medium confidence interval of 0.4. The STRING network is composed of the functional protein associations based on genomic context, high-throughput experiments, co-expression, and scientific reports. Functional enrichments in the network were identified and a subset of the identified biological processes and pathways that are relevant for metastasis were selected for visualization. The nodes in the network are colored according to their membership in each of the identified pathways.

## Acknowledgments

S.K.P. acknowledges the support of the SLAS Graduate Education Fellowship Grant. N.R. acknowledges support from the National Cancer Institute (R01CA238025). I.B. acknowledges support from the National Cancer Institute (R01 CA238025), the National Institute of Biomedical Imaging and Bioengineering (2-P41-EB015871-31), and the National Institute of General Medical Sciences (DP2GM128198). The schematic in Figure 1 was partially created with BioRender.com. The authors thank Dr. Fred Miller, Dr. Sendurai A. Mani, and Dr. Li Yang for providing 4T1 family cell lines, 4T1-FOXC2-shRNA cells, and 4T1-CXCR3-shRNA cells, respectively.

