## Supporting Information for "Raman spectroscopy reveals phenotype switches in breast cancer metastasis"

The authors disclose no potential conflicts of interest.

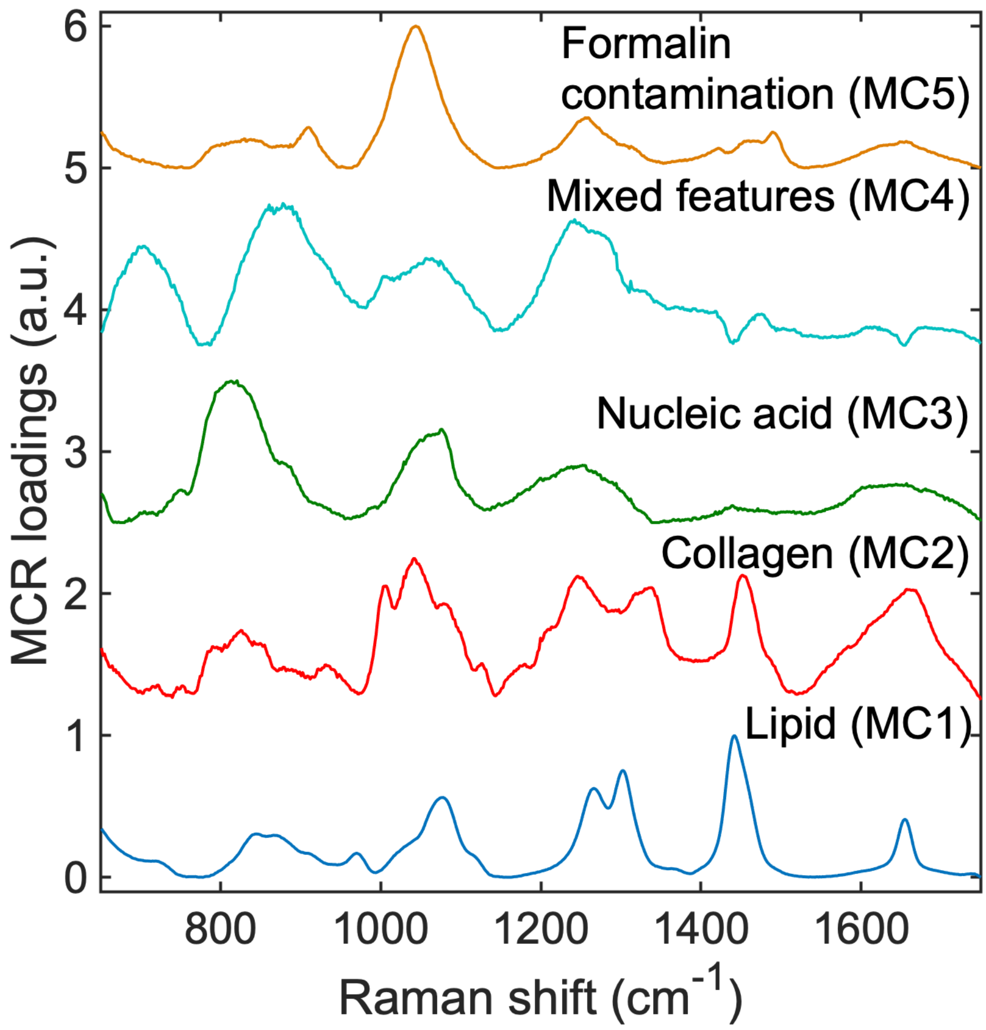


**Figure S1.** The complete set of MCR-ALS component loadings are provided for the entire Raman spectral dataset.

**Table ST1.** Table of MCR-ALS component spectral peak assignments

| Observed Raman peaks in the MCR loadings ($cm^{-1}$) | | | | | Raman band assignment from literature |
|---|---|---|---|---|---|
| **MC1** | **MC2** | **MC3** | **MC4** | **MC5** | |
| | | | 703 | | Cholesterol |
| | | 813 | | | O-P-O stretching in DNA and RNA |
| | 851 | | | | C-C stretch of proline in collagen |
| | | | 878 | | C-C stretching, hydroxyproline of collagen |
| | | | | 910 | Formalin contamination during tissue fixation |
| | 931 | | | | C-C vibration in collagen backbone |
| | 1003 | | 1003 | | Phenylalanine of collagen |
| | 1042 | | | | Proline in collagen |
| | | | | 1044 | Formalin contamination during tissue fixation |
| | | | 1063 | | Skeletal C-C stretch of lipids |
| | | 1076 | | | $PO_2^-$ symmetric stretching in DNA |
| 1078 | | | | | C-C stretch |
| | 1082 | | | | Carbohydrate residues of collagen |
| | | 1239 | 1242 | | $PO_2^-$ asymmetric stretching in DNA |
| | 1251 | | | | Amide III in collagen |
| | | | | 1256 | Formalin contamination during tissue fixation |
| 1266 | | | | | $CH_2$ in-plane deformation (Triglyceride) |
| 1302 | | | | | CH vibration (Triglyceride) |
| | 1337 | | | | $CH_3CH_2$ wagging modes of collagen |
| 1442 | | | | | $CH_2$ bending mode (Triglyceride) |
| | 1451 | | | | $CH_2$ bending mode in collagen |
| | | | | 1491 | Formalin contamination during tissue fixation |
| 1654 | | | | | C=C lipid stretch |
| | 1657 | | | | α-helical structure of amide I in collagen |

**Table ST2.** Assignment for the top spectral predictors derived from random forest analysis

| **Observed Raman peaks** | **Raman band assignment from literature** |
|---|---|
| 670 | C-S stretching mode of cystine (collagen type I) |
| 862 | Tyrosine (collagen) |
| 962 | Symmetric stretching vibration of $PO_4^{3-}$ |
| 1042 | Proline (collagen) |
| 1098 | C-N stretch (lipid) |
| 1213 | C-N stretch |
| 1254 | Amide III |
| 1330 | Collagen |
| 1372 | Ring breathing modes of DNA/RNA bases |
| 1432 | $CH_2$ deformation (lipid) |
| 1462 | $CH_2$/$CH_3$ deformation (lipid and collagen) |
| 1566 | Tryptophan |
| 1664 | Amide I (collagen) |
